# Tibanna: software for scalable execution of portable pipelines on the cloud

**DOI:** 10.1101/440974

**Authors:** Soohyun Lee, Jeremy Johnson, Carl Vitzthum, Koray Kırlı, Burak H. Alver, Peter J. Park

**Author notes:** Equally contributed (Soohyun Lee, Jeremy Johnson).

## Abstract

**Summary:** We introduce Tibanna, an open-source software tool for automated execution of bioinformatics pipelines on Amazon Web Services (AWS). Tibanna accepts reproducible and portable pipeline standards including Common Workflow Language (CWL), Workflow Description Language (WDL) and Docker. It adopts a strategy of isolation and optimization of individual executions, combined with a serverless scheduling approach. Pipelines are executed and monitored using local commands or the Python Application Programming Interface (API) and cloud configuration is automatically handled. Tibanna is well suited for projects with a range of computational requirements, including those with large and widely fluctuating loads. Notably, it has been used to process terabytes of data for the 4D Nucleome (4DN) Network.

**Availability:** Source code is available on GitHub at https://github.com/4dn-dcic/tibanna.

## 1 Introduction

Efficient execution of data processing and analysis pipelines is essential in many areas of research that involve large datasets. However, integration of a pipeline with local computing infrastructure is often a painstaking process. The need for such integration leads to redundant creation of seemingly identical pipelines, decreasing reproducibility and efficiency.

Cloud platforms present a number of advantages for large-scale data analysis, including scalability and efficient data sharing. However, the frequent lack of separation between workflow specifications and platform-specific configuration impedes transition to the cloud. Having to handle detailed aspects of the cloud platform renders the advantages of cloud computing impractical for many pipeline developers.

For effective separation of pipelines from platforms, pipelines should be made portable and standardized, and all platform-specific tasks must be delegated to a separate pipeline management tool. To enable this framework, there have been efforts to create a standard for portable pipelines. Common Workflow Language (CWL) (https://www.commonwl.org/) and Workflow Description Language (WDL) (https://software.broadinstitute.org/wdl/) describe the structure of a pipeline (e.g., steps, inputs and outputs), whereas Docker and Singularity (Kurtzner et al., 2017) enable generation of portable images of executable components and dependencies. Support for standard pipeline languages is becoming more widely adopted. In addition, Galaxy (Giardine et al. 2005), a GUI-based bioinformatics analysis platform, is planned to support CWL in the near future. Domain Specific Languages (DSLs) such as Nextflow (Tommaso et al., 2017) or Snakemake (Köster and Rahmann 2012) provide better expressive power than CWL, but are bound to a single pipeline manager.

At the Data Integration and Coordination Center (DCIC) for the NIH-sponsored 4D Nucleome (4DN) Network (Dekker at al. 2017), our aim is to efficiently process a large number of datasets from multiple experimental types submitted by member laboratories. We therefore require a workflow management system that supports standardized and portable pipelines (CWL/WDL) on a public platform. We chose Amazon Web Services (AWS), the most widely used commercial cloud service. Additionally, the Network needs an open-source implementation so the scientific community can easily run the same 4DN pipelines on its own data.

## 2 Results

We have developed and used Tibanna as an open-source pipeline manager and automated cloud resource allocator for running Docker-based pipelines on AWS. Tibanna integrates easily with other systems through its Python API.

### 2.1 Overview

Tibanna creates an instance of an individually pre-configured virtual machine (EC2) for each job (Figure 1). The instance autonomously fetches data and the pipeline, runs the pipeline, stores logs and output files on the cloud and finally, terminates itself. Instead of depending on a master server or queue, each job gets its own serverless scheduler comprised of AWS Lambdas coordinated by the AWS Step Function. The 4DN DCIC uses additional Lambdas to communicate with our Data Portal.

**Fig 1.**
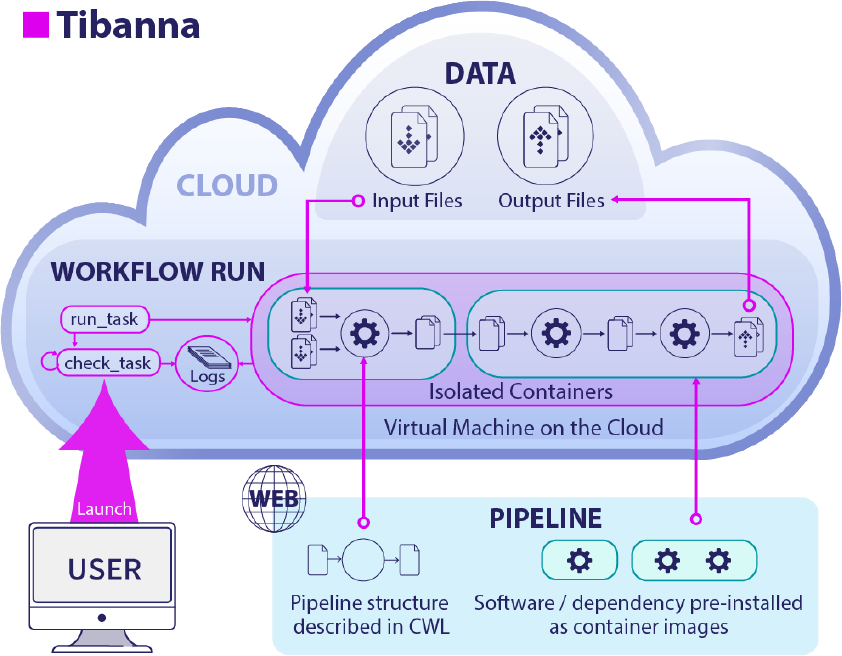
Tibanna. For each pipeline execution, Tibanna creates a workflow run by launching a customized virtual machine (EC2), runs the pipeline and destroys itself when finished. Scheduling is performed individually by serverless components.

### 2.2 Execution unit

Tibanna runs a single execution unit, and dependencies can be managed through its dependency feature. This allows one to decide and manage execution units based on resource requirements and logical grouping, and determine what intermediate files to keep for future re-processing and curation. An execution unit may be a whole pipeline, a sub-pipeline or a single step, and these units are not defined by the pipeline structure itself. We found that tools that attempt to auto-determine execution units tend to make non-optimal choices and are difficult to integrate with desired execution designs.

### 2.3 How to use Tibanna

Tibanna auto-configures cloud components and permissions with a single command. Its only requirements are to provide input data on the cloud and to prepare a publicly available pipeline in CWL or WDL, along with a public Docker image. Each job is described as a JSON file or a Python dictionary object, specifying the pipeline, input files and parameters, and where the output should be collected. Submission and monitoring of jobs is performed locally from the command line or using the Python API. Logs can also be retrieved similarly or through the AWS Web Console. A user with a private key file can securely connect to running instances for more detailed monitoring. An easy-to-follow documentation comes with the software for more details.

### 2.4 Execution example

In addition to running pipelines for 4DN, Tibanna has been used by external users to call transposon insertions from 1,000 ~30X whole genome sequencing data sets. 1,000 jobs were simultaneously executed, each running for 4-8 hours on a spot instance with 16 cores and 32GB memory and 250GB disk size, costing about $1.6 per run.

### 2.5 Comparison to other tools

Tibanna offers several advantages compared to existing workflow management tools (Supplementary Table 1), specifically for our need to automatically execute and monitor various portable pipelines in integration with a data portal.

Local CWL executors (e.g. Cwltool, Rabix (Kaushik 2016) and CWL-Airflow (Kotliar et al. 2017)) do not perform resource allocation. Snake-make recently started supporting local CWL executions with cloud storage but also does not handle resource management on AWS.

Tools that perform cloud resource allocation uses one of the three approaches; AWS Batch, a resizable cluster, or individual customization.

#### (1) AWS Batch

AWS Batch (https://aws.amazon.com/batch/) is an AWS service to run batch jobs, but it is not easy to be used directly with CWL/Docker, often exhibits a mysteriously long wait time, and lacks many functionalities. Examples of tools based on AWS Batch include Nextflow, Funnel (https://ohsu-comp-bio.github.io/funnel/) and Cromwell. Nextflow supports Docker, but using CWL requires conversion to the Nextflow format and the converter is not functional at this point.

Cromwell (https://software.broadinstitute.org/wdl/) was originally developed to run WDL on Google Cloud Platform (GCP), although it now supports CWL and AWS through AWS Batch. However, it leaves all the cloud configuration to the user, and due to the lack of an API, a large-scale batch run requires a cumbersome generation of many input JSON files and manual tracing of job IDs for proper output matching. Monitoring functionality that is critical for large-scale projects is not provided by either Cromwell or AWS Batch. Funnel also requires users to build a custom Amazon Machine Image (AMI).

#### (2) Resizable cluster

StarCluster (http://star.mit.edu/cluster/docs/latest), an older tool without CWL/Docker support, and Toil (Vivian et al., 2017) work by creating a resizable cluster of EC2 instances of the same CPU and memory and an additional master instance. They require extra work for configuring and managing clusters. Nextflow also uses an auto-scalable cluster as an alternative. Arvados (https://arvados.org/) supports CWL and Docker and involves a cluster of dynamically allocated cloud instances where jobs are dispatched by SLURM. However, the initial setup process is quite complicated. Arvados also uses its own storage organization which required additional development to access and is not compatible with the file organizations used by our portal.

#### (3) Individual customization

Tibanna instead creates a specific compute environment for an individual execution at launch, and removes it when the job is done. This approach is also adopted by Seven Bridges (https://www.sevenbridges.com/) and DNANexus (https://www.dnanexus.com/), which, as commercial portal services, also offer full automation but charge extra fees and do not easily integrate with other systems.

At 4DN DCIC, we receive data sets that vary in input file size, with vastly different memory and storage requirements. For example, the contact matrix generation sub-workflow of the 4DN Hi-C data processing pipeline involves in-memory generation of a matrix that represents nonzero interactions of all genomic locations. As the size of input datasets (i.e., reads representing contacts) varies >10-fold (up to 76Gb), memory requirement also varies substantially, thus necessitating machines with different memory sizes for efficient computation. For this level of variability at the execution level, individual customization makes better use of the elastic compute environment of the cloud compared to cluster allocation.

## 3 Conclusion

Tibanna is a pipeline management system for the AWS cloud. Tibanna has been used to process 4DN data. We believe Tibanna will be useful for many other AWS projects.

## Acknowledgements

We thank the members of the 4DN DCIC, the Park lab and the Avillach lab at the Department of Biomedical Informatics at Harvard Medical School for valuable feedback and the E. Alice Lee lab at Boston Children’s Hospital for beta-testing Tibanna. We are especially thankful to Andy Schroeder, Peter Kerpedjiev, Martin Owens and Chuck McCallum for code review/contribution, Shannon Ehmsen for preparing the figure, and Alon Galor, Dhawal Jain, Su Wang, Geoff Nelson and Marc Rubenfield (Veritas Genetics) for comments on the manuscript.

---

**Supplementary Table 1.** Comparison of cloud-friendly pipeline executors. Cloud-friendly pipeline executors are compared in various aspects, focused on compatibility with portable languages and degree of support for AWS cloud.

|  | Tibanna | Seven Bridges | DNA Nexus | Nextflow | Toil | Arvados | Cromwell | Snakemake |
| --- | --- | --- | --- | --- | --- | --- | --- | --- |
| **Common Workflow Language (CWL) support** | yes | yes | no | no | yes | yes | yes | yes |
| **Workflow Description Language (WDL) support** | yes | no | yes | no | yes | no | yes | no |
| **computing on Amazon Web Services (AWS)** | yes | yes | yes | yes | yes | yes | yes | no |
| **computing on Google Cloud Platform (GCP)** | no | yes | no | yes | yes | yes | yes | yes |
| **easy to integrate with user's own cloud data storage system** | yes | moderate | no | yes | yes | no | yes | yes |
| **free and open source** | yes | no | no | yes | yes | yes | yes | yes |
| **Graphic User Interface (GUI)** | via AWS | yes | yes | no | no | yes | no | no |
| **Command-line interface (CLI)** | yes | yes | yes | yes | yes | yes | yes | yes |
| **Python Application Programming Interface (API)** | yes | yes | yes | no | yes | yes | no | yes |
| **easy setup on AWS cloud** | yes | yes | yes | no | no | no | no | - |
| **cloud resource allocation approach on AWS** | individual | individual | individual | cluster<br>AWS Batch | cluster | cluster | AWS Batch | - |
| **master server required** | no | no | no | yes | yes | yes | no | no |
| **Comes with its own portal service** | no | yes | yes | no | no | yes | no | no |

